# Pathology and immunity after SARS-CoV-2 infection in male ferrets is affected by age and inoculation route

**DOI:** 10.1101/2021.06.30.450298

**Authors:** Koen van de Ven, Harry van Dijken, Lisa Wijsman, Angéla Gomersbach, Tanja Schouten, Jolanda Kool, Stefanie Lenz, Paul Roholl, Adam Meijer, Puck van Kasteren, Jørgen de Jonge

## Abstract

Improving COVID-19 intervention strategies partly relies on animal models to study SARS-CoV-2 disease and immunity. In our pursuit to establish a model for severe COVID-19, we inoculated young and adult male ferrets intranasally or intratracheally with SARS-CoV-2. Intranasal inoculation established an infection in all ferrets, with viral dissemination into the brain and gut. Upon intratracheal inoculation only adult ferrets became infected. However, neither inoculation route induced observable COVID-19 symptoms. Despite this, a persistent inflammation in the nose was prominent in especially young ferrets and follicular hyperplasia in the bronchi developed 21 days post infection. These effects -if sustained- might resemble long-COVID. Respiratory and systemic cellular responses and antibody responses were induced only in animals with an established infection. We conclude that intranasally-infected ferrets resemble asymptomatic COVID-19 and possibly aspects of long-COVID. Combined with the increasing portfolio to measure adaptive immunity, ferrets are a relevant model for SARS-CoV-2 vaccine research.

## Introduction

Severe acute respiratory syndrome coronavirus 2 (SARS-CoV-2) was first detected in patients near the end of 2019 and soon started a new pandemic ^1^. As an intervention, effective SARS-CoV-2 vaccines were rapidly developed and implemented. Although many of these vaccines have proven to be effective in limiting mortality and morbidity ^2^, it remains a point of concern that SARS-CoV-2 mutants might escape vaccine-induced immunity ^3^. Additionally, much remains unknown about the pathology and long-term effects of SARS-CoV-2 infection and protective immunity. Animal models are essential for investigating these issues and the scientific community has made unprecedented advances in their development since the outbreak ^4, 5^. However, there are still some remaining knowledge gaps that limit the evaluation of outstanding questions.

SARS-CoV-2 spreads through direct contact, aerosols, or droplets ^6–9^. Viral RNA has been detected in stool samples of infected individuals ^10–12^ and this has raised the question whether the fecal-oral route could also facilitate SARS-CoV-2 transmission ^13^. The virus primarily replicates in respiratory tissue, but viral RNA has been detected in many other tissues including the brain, gut, heart, and endothelial lining of the vascular system ^12, 14, 15^. SARS-CoV-2 infection usually induces mild disease (COVID-19), with symptoms limited to fever, dry cough, anosmia/ageusia, myalgia, fatigue, dyspnea, sputum production, headache, and occasionally diarrhea (reviewed in ^16^). Fatal cases are marked by respiratory failure, arrhythmia, shock, and acute respiratory distress syndrome ^15, 17, 18^. Mortality significantly increases with age and certain comorbidities such as cardiovascular disease and diabetes ^15, 17, 19–21^. Despite similar infection rates between men and women, men are more likely to succumb to infection ^17, 19, 20^. Some individuals additionally suffer from ‘long-COVID’ (post-acute COVID-19 syndrome) where they experience persisting symptoms long after initial SARS-CoV-2 infection ^22^.

As the ferret (*Mustela putorius furo*) is considered the best small animal model for respiratory disease caused by influenza virus ^23, 24^, multiple groups have investigated if ferrets are also suited to model COVID-19 ^25–32^. From these studies, we know that SARS-CoV-2 efficiently replicates in the upper respiratory tract (URT) of ferrets upon intranasal (i.n.) inoculation, but replication in the lower respiratory tract (LRT) is limited ^25^. There are however still many unknowns regarding the ferret model.

Here, we report our efforts to investigate the influence of both age and inoculation route on SARS-CoV-2 infection in the ferret model. As intratracheal (i.t.) inoculation with influenza virus induces more severe disease in ferrets ^33–35^, we wondered if we could model severe COVID-19 in ferrets by i.t. inoculation with SARS-CoV-2. In addition, as advancing age is a risk factor for the development of severe disease ^17, 19, 20^, we investigated the role of age by infecting both young and adult ferrets. We found that SARS-CoV-2 infection was more efficient via the i.n. route and that i.t. inoculation and increased age did not result in more severe disease. Regardless of age and inoculation route, no clinical disease was observed. Despite the absence of symptoms, we did find pathological aberrations in the nose and lungs that were more prominent in young ferrets and seemed to increase with time, potentially reflecting long-COVID in humans. We also report that humoral and cellular immune responses appear to depend on sufficient viral load during the acute phase of the infection. Together, our findings indicate that while ferrets might not be suited to study severe COVID-19, they can be used to model viral replication, adaptive immune responses, asymptomatic COVID-19 and possibly long-COVID.

## Results

### Study outline

In this study we assessed the role of age and infection route on SARS-CoV-2 disease and immunity in ferrets, in an attempt to model the (severe) COVID-19 observed in humans. We infected young (9-10 months) and adult (36-48 months) male ferrets with SARS-CoV-2 through either intranasal (i.n.) or intratracheal (i.t.) inoculation (Fig. 1a). To show that adult ferrets differed from young ferrets immunologically, we performed baseline whole blood trucounts. Compared to young ferrets, B and T cell numbers were lower in adult ferrets (Fig. 1b and Supplemental Fig. 1), which has also been described for older humans ^36^. Ferrets were then inoculated with a dose of 10^7^ TCID_50_ as others have reported that a low dose is insufficient to establish infection ^32^. Mock-infected young and adult ferrets were inoculated i.n. with PBS. On 5, 14, and 21 days post infection (dpi), three animals per group were euthanized to study viral replication, pathology, and immune responses. Due to a limited supply of ferrets, no adult i.n. inoculated animals were euthanized on 14 dpi.

**Figure 1:**
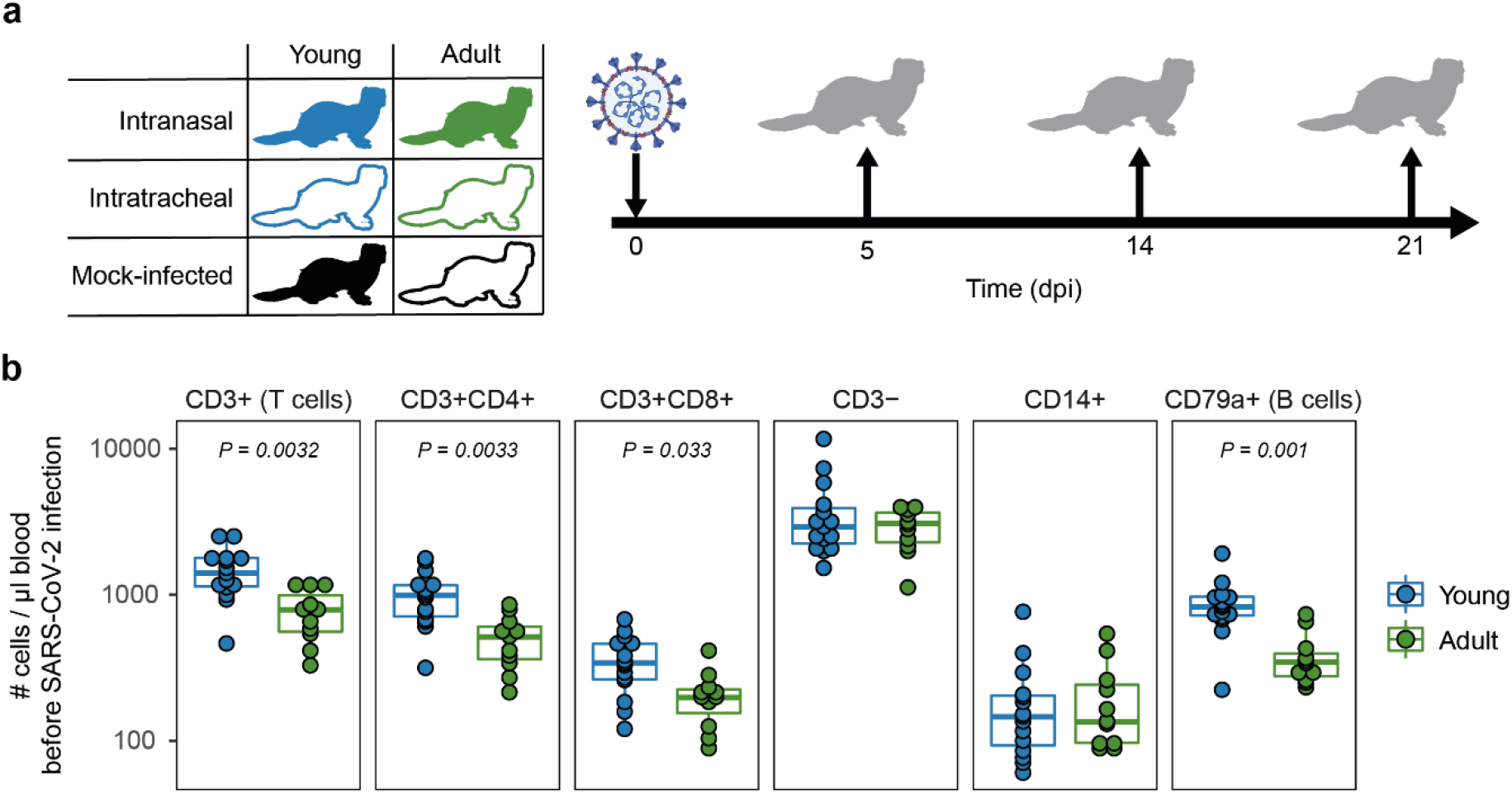
study outline and immunological profile of young and adult ferrets. **A**) Young and adult ferrets were inoculated intranasally or intratracheally with 10^7^ TCID_50_ SARS-CoV-2 on day 0. On 5, 14 and 21 days post infection (dpi), 3 ferrets were euthanized per group. Mock-infected ferrets were euthanized on 14 dpi (young) and 21 dpi (adult). No intranasally inoculated adult ferrets were euthanized at 14 dpi due to limited availability of adult ferrets. **B**) Cell counts in blood of uninfected young and adult ferrets. N = 15 for young animals and n = 11 for adult animals. Boxplots depict the median with the first and third quartiles. Differences between young and adult ferrets were tested via a two-sample Wilcoxon test and corrected for multiple testing using the Holm method.

### Viral load is higher after i.n. inoculation

After i.n. SARS-CoV-2 infection, viral RNA was high in nose, throat, and rectal swabs, with no difference between ages (Fig. 2a). In contrast, i.t. inoculation led to reduced viral loads and was clearly influenced by age as less viral RNA was measured in young animals. Viral RNA detected by RT-qPCR does not necessarily equal the presence of infectious virus, so we determined the amount of replication-competent virus in the nose and throat by TCID_50_-assay. Although viral RNA could be detected by RT-qPCR as late as 21 dpi (Fig. 2a), infectious virus was no longer detectable by 9 dpi (Fig. 2b). I.n. infection resulted in the highest viral titer in nose and throat. Viral titers were lower for i.t. infected ferrets, especially for young ferrets where almost all samples were below the detection limit.

**Figure 2:**
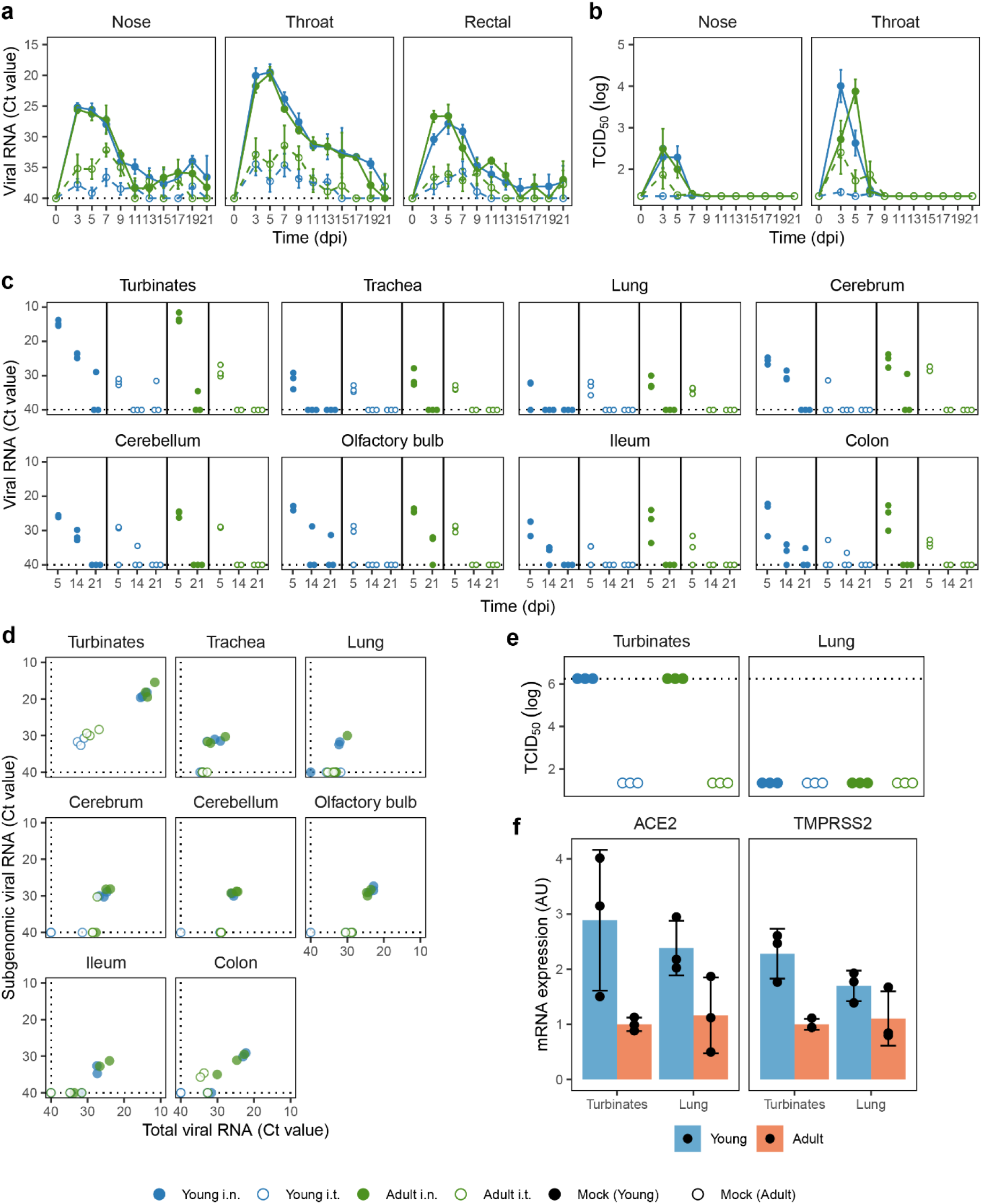
Viral load in swabs and tissues of SARS-CoV-2 infected ferrets. **A, B**) Viral load measured by RT-qPCR (a) and TCID_50_-assay (b) in swabs collected from SARS-CoV-2 infected ferrets on various days post infection (dpi). **C**) Viral RNA in tissues at 5, 14 and 21 dpi measured by RT-qPCR. **D**) Total viral RNA on 5 dpi as depicted in (c) plotted against subgenomic viral RNA. **E**) Infectious virus detected by TCID_50_-assay in nasal turbinates and lung 5 dpi. **F**) Expression of SARS-CoV-2 receptor ACE2 and TMPRSS2 protease in nasal turbinate and lung tissue of placebo animals, plotted as arbitrary units (AU). In panels a, c and d, RT-qPCR negative specimens were set to a Ct-value of 40 for visualization purposes, which is depicted by dotted lines. In panel e, dotted lines depict the highest dilution tested. For a, b: n = 3-9. For c-f: with exception of ‘Adult i.t.’ on 14 dpi (n = 2), all groups are n = 3.

Next, we measured viral RNA in various tissues on 5, 14, and 21 dpi. Similar to the swabs, viral RNA was higher in almost all tissues of i.n. inoculated ferrets at 5 dpi, independent of age (Fig. 2c). For i.n. infected ferrets, viral RNA was high in the nasal turbinates and low in the lung. Contrary to our expectations, i.t. inoculation did not lead to more viral RNA in the LRT (lungs and trachea). Viral RNA was also detected in the gut (ileum and colon) and disseminated from the initial site of infection to the olfactory bulb, cerebrum and even into the cerebellum. In these tissues, more viral RNA was detected in ferrets infected i.n. From day 5 onwards, viral RNA declined and was only sporadically detectable at 21 dpi.

To test whether the detected viral RNA was a result from an active infection at the site of sampling, we performed both a TCID_50_-assay and an RT-qPCR specific for viral subgenomic mRNA. The presence of viral subgenomic mRNA is indicative of previous or current viral transcription in infected cells and was detected in most tissues 5 dpi (Fig. 2d), but declined afterwards. By 14 dpi viral subgenomic mRNA was only detected in the nasal turbinates of young i.n. inoculated ferrets and at 21 dpi all tissues tested negative (data not shown). Interestingly, while the presence of subgenomic mRNA indicates that SARS-CoV-2 infected multiple tissues 5 dpi, this did not result in the production of detectable infectious viral particles. Analysis by TCID_50_ showed that infectious virus was only produced in the nasal turbinates at 5 dpi, with no detectable infectious virus in the lung, gut or brain at 5 dpi or later timepoints (Fig. 2e and data not shown). Due to limited tissue availability, viral TCID_50_-titers in nasal turbinates were not tested on 14 and 21 dpi.

Infection of cells by SARS-CoV-2 is dependent on the expression of the binding receptor ACE2 (Angiotensin-converting enzyme 2) and the fusion priming protease TMPRSS2 (Transmembrane protease, serine 2) ^1, 37, 38^. In order to determine if the reduced replication of SARS-CoV-2 in the LRT of young animals could be explained by lower ACE2 and TMPRSS2 expression, we quantified ACE2 and TMPRSS2 mRNA by RT-qPCR in young and adult placebo animals. ACE2 and TMPRSS2 expression was however similar between nasal turbinates and lung tissue, indicating that the absence of SARS-CoV-2 replication in the LRT was not due to reduced expression of ACE2 or TMPRSS2 (Fig. 2f). Adult animals did seem to express lower levels of ACE2 and TMPRSS2, but this did not negatively impact viral load (Fig. 2a, b).

### Pathology in the URT is higher for i.n. infected animals

Despite active SARS-CoV-2 replication, ferrets did not display any overt clinical signs of disease. Compared to mock-infected animals, SARS-CoV-2-infected animals did not lose more weight and did not experience fever (Supplemental Fig. 2ab), nor were alterations in respiratory function and physical activity observed. Despite the absence of clinical disease, infection with SARS-CoV-2 did result in pathological aberrations in the respiratory tract.

In the nasal turbinates, pathological aberrations were restricted to the respiratory epithelial lining of the naso- and maxilloturbinates (Fig. 3a). The sub- and intraepithelial inflammation resulted in thickened respiratory mucosa and an exudate of polymorphonuclear cells, lymphocytes and some macrophages. Damage to the epithelial lining was characterized by hypertrophy, hyperplasia, and squamous metaplasia and sporadically the epithelium was absent. Hypertrophy of goblet cells was also present and haemorrhage was sometimes observed. The different pathological facets were categorized into inflammation and damage, for which an end-score was determined as previously described (Fig. 3b, c and Supplemental fig. 3) ^39^.

**Figure 3:**
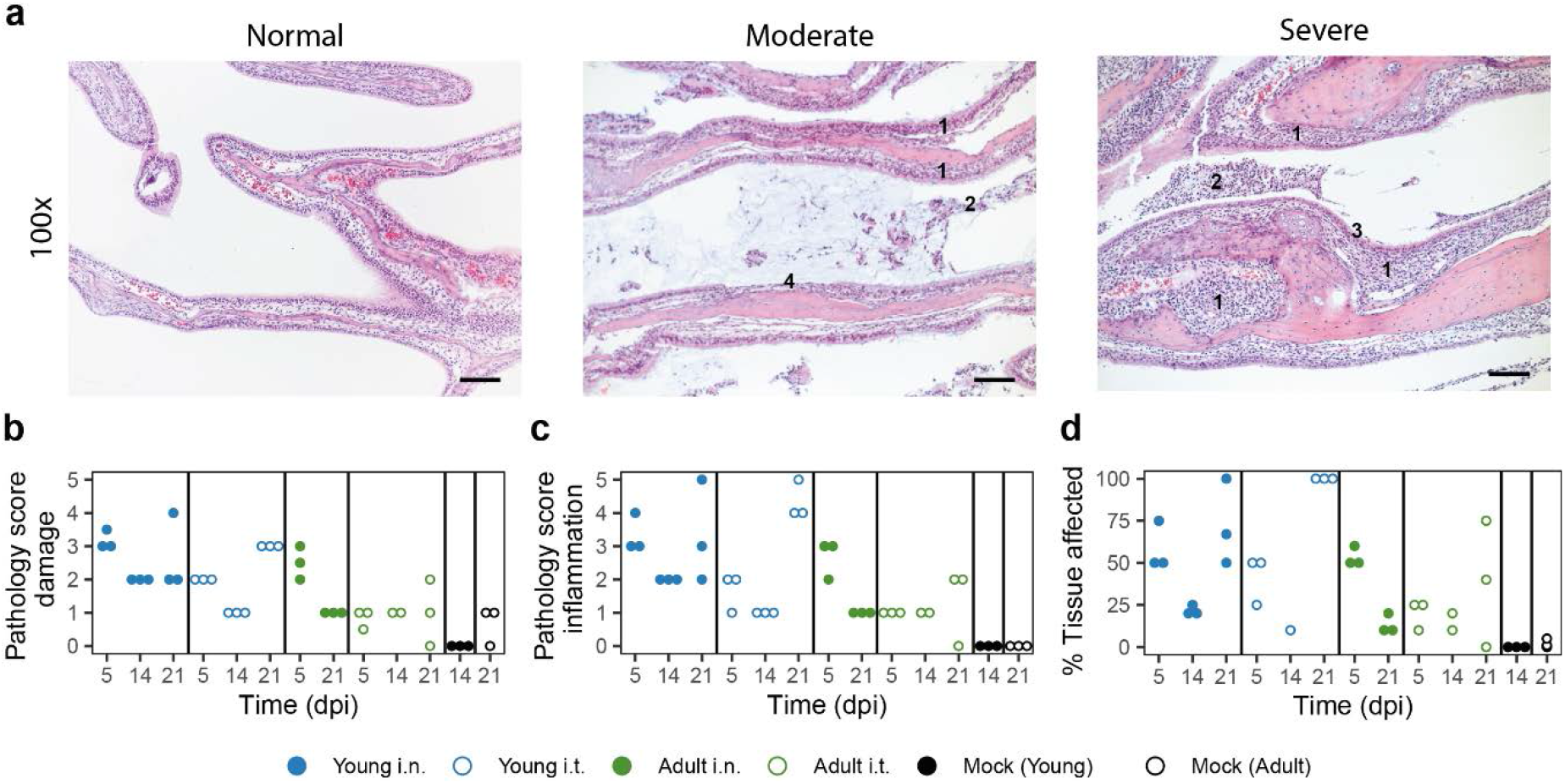
SARS-CoV-2 infection induces more severe URT pathology in young ferrets. **A**) Representative haematoxylin and eosin stained sections of normal, moderate (score 2) and severe (score 4) affected naso- and maxilloturbinates at 100x magnification. SARS-CoV-2 infection induced infiltration of polymorphonuclear cells into the lamina propria and submucosa of the respiratory epithelium, resulting in thickened respiratory mucosa (indicated with ‘1’). This was accompanied by exudate containing mucus (moderate) and many polymorphonuclear cells, lymphocytes and some macrophages (2). The infection caused hyperplasia/hypertrophy of the epithelial cells (3) and sometimes the epithelial lining was absent (4). Bars represent 50μm. **B-D**) Scoring by parameters related to epithelial damage (b), inflammation (c) and percentage of tissue involved in pathologic changes (d). The infection-induced pathology was scored on a scale of 0–5 based on the parameters described in the materials and methods. With exception of ‘Adult i.t.’ on 14 dpi (n = 2), all groups are n = 3 and each dot represents an animal.

In general, the severity of pathology was dependent on age and infection route, although the age-correlation was opposite to our expectations. In the acute phase (5 dpi), i.n. inoculated ferrets displayed more pathological aberrations in the nasal turbinates, which partly resolved by 14 dpi (Fig. 3b, c). Strikingly, inflammation and to a lesser extent epithelial damage increased again by 21 dpi in young animals, but was less pronounced in adult ferrets. Notably, this effect was stronger in i.t. infected young ferrets than in i.n. infected young ferrets and involved 100% of the epithelium of the naso- en maxilloturbinates (Fig. 3d). Pathology in the nose thus increased in young animals even after the infection had been cleared.

### Pathology in the LRT is not affected by inoculation route

On a macroscopic level, the lungs of SARS-CoV-2 infected ferrets did not show aberrations in the acute phase (5 dpi), apart from a few darker patches. However, red opalescent coloring along the bronchus and as isolated patches started to appear at 14 and 21 dpi. During the course of this study, there was no increase in lung weight, indicating absence of serious edema (Supplemental Fig. 2c). Microscopically, slight peribronchi(oli)tis – characterized by the presence of infiltrating cells in the sub-mucosa along the bronchi – was observed 5 dpi (Fig. 4a). The bronchus epithelium showed some reaction in the form of mild to minor hyperplasia, sometimes visible as repeated epithelial bumps. Strikingly, hyperplasia of the Bronchus-Associated Lymphoid Tissue (BALT) developed in 1 out of 3 i.n. and in 2 out of 3 i.t. infected young ferrets by 21 dpi, but was absent in adult ferrets. Mild to strong follicular hyperplasia was located at the first branches of the bronchus and in the more severe cases extended to the smaller bronchiole (Fig. 4a). Obstruction of bronchioli occurred regularly due to compression by the hyperplastic follicles. These consisted of activated lymphocytes, which penetrated through the muscle layer of the bronch(iol)us into the lamina propria. In addition, albeit low in number, local patches of mild to moderate desquamative interstitial pneumonia had developed. The pathology score and percentage affected lung parenchyma increased from 5 to 21 dpi, mostly in in the young ferrets (Fig. 4b-c and Supplemental Fig. 4). Of note, young animals that were investigated 21 dpi had tested positive for antibodies against NL-63 and enteric and systemic corona viruses prior to SARS-CoV-2 infection (Supplemental table 1). This combination was absent in adult ferrets, complicating the interpretation of these results.

**Figure 4:**
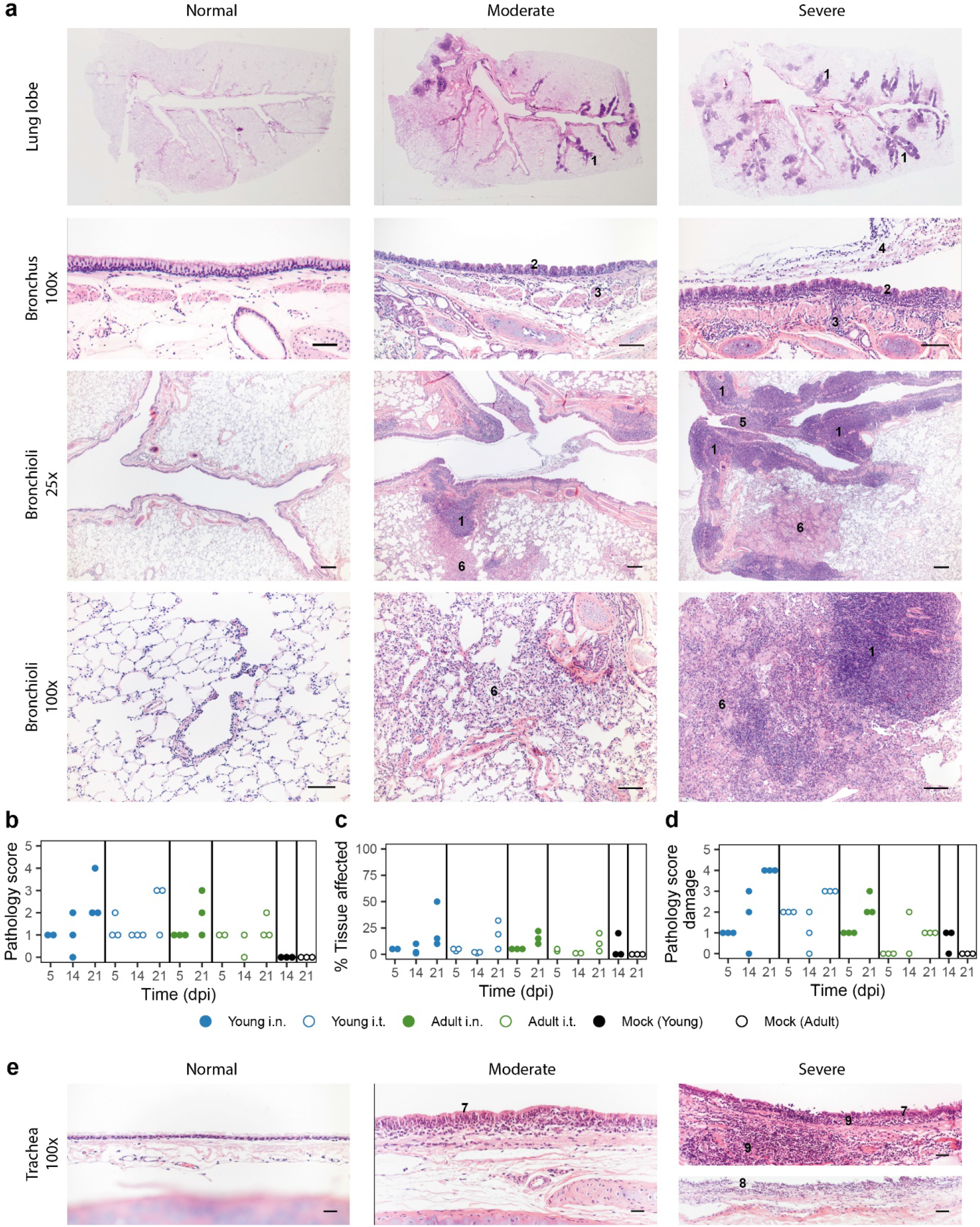
Pathology aspects of lung and trachea. Representative haematoxylin and eosin stained sections of normal, moderately and severely affected lung tissue (a) at overview, 25x and 100x magnification and trachea (e) at 100x magnification. **A**) Lungs of SARS-CoV-2 infected animals with a clear hyperplasia of the Bronchus-Associated Lymphoid Tissue (BALT), consisting of activated lymphocytes that penetrate through the muscle layer of the bronch(iol)us into the lamina propria and submucosa (indicated in figure with ‘1’). Hyperplasia of bronchial epithelium visible as repeated bumps (2), cellular infiltrate into the submucosa (3) and exudate in the lumen (4) were also observed. The presence of hyper-plastic follicles led to obstruction of bronchiole (5). Some animals displayed desquamative interstitial pneumonia, with pulmonary macrophages in the alveoli and to a minor extent, lymphocytes and plasma cells (6). The alveolar epithelium (pneumocytes) exhibited squamous-like characteristics. **B-D**) Pathology was summarized in an overall score ranging from 0-5 based on the parameters described in the materials and methods. Pathology score (b) and % of tissue affected (c) for lung tissue and pathology score for trachea (d). With exception of ‘Adult i.t.’ on 14 dpi (n = 2), all groups are n = 3. **E**) Pathological aberrations in the trachea consisted of hyperplasia (7), damage and pseudo-squamous characteristics of the epithelium (8) and infiltration of inflammatory cells into the submucosa and epithelium of the trachea (9). Bars represent 100μm (a) and 50μm (e).

In concordance with our findings in the lung and nose, pathology in the trachea increased by 21 dpi (Fig. 4d, Supplemental Fig. 5). The epithelium of the trachea in all young ferrets and in the i.t. adult group was often hyperplastic and sometimes showed serious damage and pseudo squamous characteristics 21 dpi (Fig. 4e). Inflammatory cells infiltrated the submucosa and epithelium, most prominently in young ferrets. Since SARS-CoV-2 viral RNA was also detected in the gut, histopathological analysis of the ileum and colon was performed, but no deviations were observed. In conclusion, independent of inoculation route, young animals displayed more severe pathology than adult ferrets in both the URT and LRT, but it is uncertain whether the increase until 21 dpi is due to age or prior exposure to other coronaviruses.

**Figure 5:**
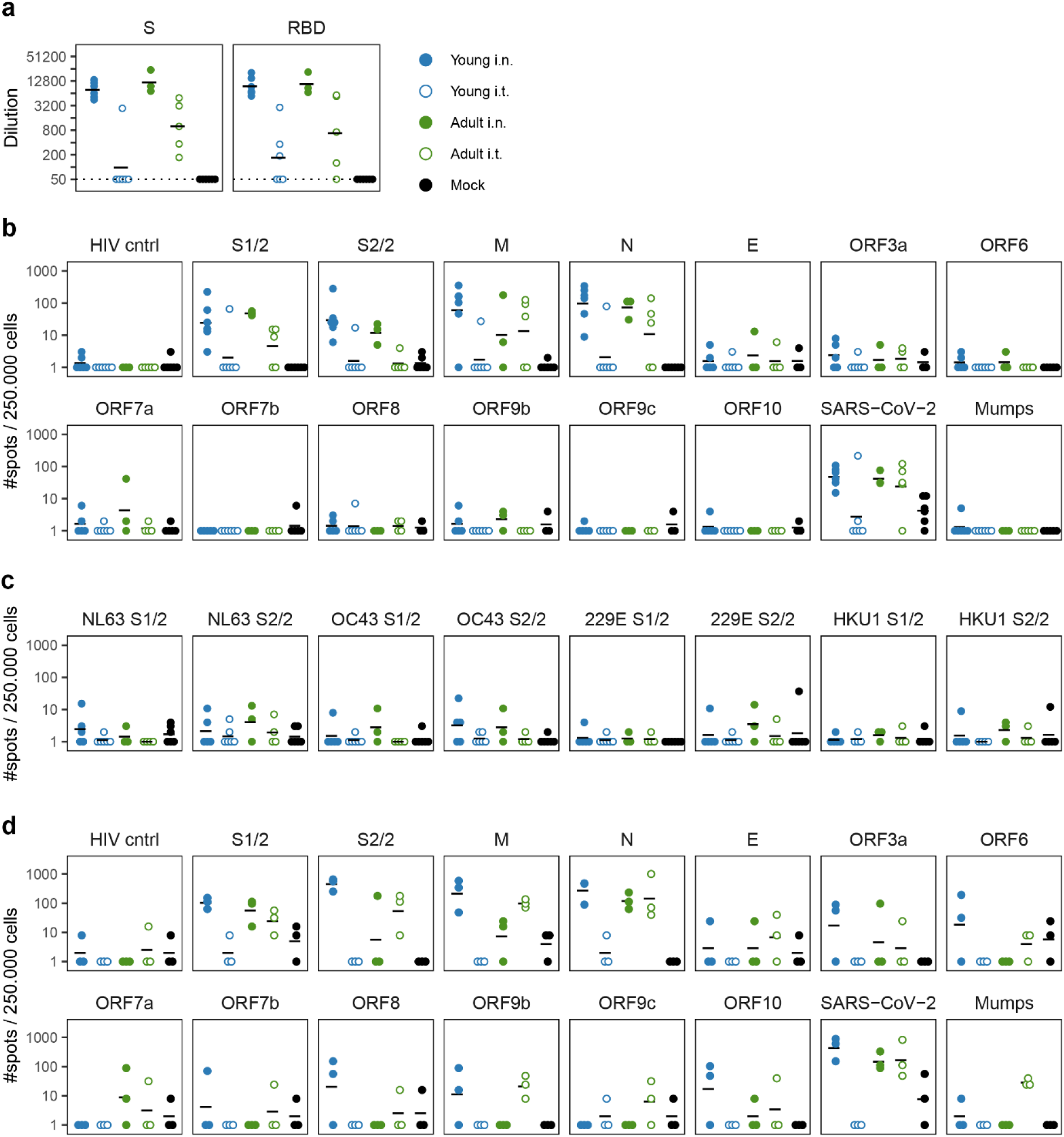
Cellular and humoral responses in SARS-CoV-2 infected ferrets are affected by inoculation route. **A**) Humoral responses detected by ELISA against whole spike (S) and the receptor binding domain (RBD) region of spike in serum 14/15 dpi. Responses are depicted as the (modelled) dilution at which the ELISA curve drops below background (mean + 3x SD of SARS-CoV-2 naïve animals at 200x dilution). The dotted line indicates the lowest dilution tested and negative samples were set to half that dilution for visualization purposes. **B-D**) Cellular responses in PBMCs (b, c) and lung-derived lymphocytes (d) as determined by IFNγ-ELISpot. Cells were stimulated with SARS-CoV-2 peptide pools or live virus. HIV cntrl and Mumps are negative controls for peptide pool and virus stimulations respectively. Data shown here were corrected for medium background and were set to a minimum of 1 spot for visualization on a log-scale. **B**) PBMCs isolated 14 or 15 days post infection (dpi). **C**) Frozen PBMCs isolated 14 or 21 dpi were thawed and stimulated with peptide pools of endemic human coronaviruses. **D**) Responses by lung-derived lymphocytes at 21 dpi. For b, c and d, samples were collected 14, 15 or 21 dpi, but visualized as a single group. N = 3-6 for panels a-c and n = 3 for panel d. Black lines indicate (geometric) mean per group.

### Cellular and humoral Immunity

As several ferrets used in this study displayed antibody responses against ferret corona viruses and NL-63 prior to the start of the study (Supplemental table 1), we wondered if cross-reactive responses to SARS-CoV-2 were present in these animals. We measured antibody responses against SARS-CoV-2 spike protein (S) and its receptor binding domain (RBD) by ELISA. Cellar responses were measured by IFNγ-ELISpot in which PBMCs were stimulated with live SARS-CoV-2 or overlapping peptide pools of the S, membrane (M) and nucleoprotein (N) proteins. In five animals the antibody response against SARS-CoV-2 was just above background prior to infection (Supplemental Fig. 6a). Additionally, another animal in the young i.n. infected group clearly responded against an overlapping peptide pool of SARS-CoV-2 spike protein in ELISpot on two separate time-points before infection (Supplemental Fig. 6b). This animal tested positive for antibodies against ferret corona and NL63, but not against SARS-CoV-2 (Supplemental table 1 and Supplemental Fig. 6a), suggesting that there might have been a cross-reactive T cell response. Interestingly, the animal displayed similar viral kinetics as other animals of the same treatment group, but it displayed more severe pathology 21 dpi.

After SARS-CoV-2 infection, i.n. inoculated ferrets displayed high antibody responses against both S and RBD with no evident differences between young and adult animals 14 dpi (Fig. 5a). In contrast, antibody titers in i.t. infected animals were lower with a clear distinction between ages. More adult (4/5) than young (1/6) i.t. inoculated ferrets displayed responses against S-protein after infection. Similar findings were obtained for humoral responses against RBD.

Next, we investigated cellular responses at 14 dpi. PBMCs mainly responded against the larger peptide pools of S (S1 & S2), M and N (Fig. 5b). We additionally measured responses against envelope (E) and accessory proteins (ORF3a-ORF10), but responses against these peptide pools were marginal and did not differ from naïve animals. Consistent with the low viral load, only 1/6 young i.t. infected animals responded to any SARS-CoV-2 stimulus, indicating that almost no cellular immunity was established in this group by 14 dpi. For the other groups, no clear differences between infection routes or age were observed. The responses 21 dpi were similar to those of 14 dpi (Supplemental Fig. 7a). As others have reported the presence of cross-reactive T cell responses in humans ^40–43^, we also tested if PBMCs from SARS-CoV-2 infected ferrets would respond upon stimulation with spike peptide pools of coronaviruses NL63, OC43, 229E and HKU1. Some animals displayed minor responses, although they were only slightly higher than mock-infected animals (Fig. 5c).

We additionally investigated cellular responses in the lung. To reduce contamination of the lung lymphocytes with circulating lymphocytes, lungs were perfused with saline on 14 and 21 dpi. In comparison to PBMCs, responses against the smaller SARS-CoV-2 peptide pools seemed more prominent in lung-derived lymphocytes of young i.n. infected animals. In this group, two out of three ferrets showed responses against ORF3a, ORF6, ORF8, ORF9b and ORF10 peptide pools at 21 dpi (Fig. 5d). Like in PBMC, cellular responses were absent in the lung of i.t. infected young ferrets. Of note, responses in the lungs of i.t. infected old animals differed between day 14 and 21 post infection, with responses being higher overall on 21 dpi (Fig. 5d and Supplemental Fig. 7b). However, the group size (n = 2-3) was too small for reliable statistical testing.

## Discussion

In this study we utilized a male ferret model to assess the influence of age and infection route on SARS-CoV-2 disease and immunity. Intranasal inoculation was more efficient in establishing an infection compared to intratracheal administration. This was especially the case for young ferrets, in which virus hardly replicated after i.t. inoculation. In contrast, disease and pathology were not increased in older animals or upon intratracheal inoculation.

Despite a productive SARS-CoV-2 infection, ferrets did not develop the symptomatic disease that is reported for humans (reviewed in ^16^). A slightly more watery defecation was observed in some animals towards the end of the study, but occurrences were too few to confidently attribute this to SARS-CoV-2 infection. We did however detect SARS-CoV-2 genomic RNA and subgenomic mRNA in rectal swabs and gut tissues at multiple time-points. Subgenomic SARS-CoV-2 mRNA is indicative of viral infection, which suggests that SARS-CoV-2 is indeed enterotropic in ferrets, similar to what has been reported for humans ^14, 44–47^. In addition, the presence of viral genomic RNA and subgenomic mRNA in the olfactory bulb, cerebrum and cerebellum of ferrets is in line with reports that SARS-CoV-2 can infect the CNS of patients ^14, 48, 49^. Infection of the olfactory bulb might explain the anosmia often found in COVID-19 patients. It is however important to note that while the presence of subgenomic mRNA indicates infection of various tissues, there was a lack of infectious virus as measured by TCID_50_-assay. This raises the question whether the production of infectious virus in these tissues is completely absent or only happened within the first few days after inoculation.

The lack of clinical disease in this study is in line with previous SARS-CoV-2 ferret studies. In those studies bodyweight did not decrease after infection, although reduced activity was observed in some instances ^25, 29, 32^. Fever was observed in some studies ^25, 27^, but not in others ^29, 32^. Ferrets thus do not model the severe aspects of COVID-19. Despite this knowledge, we initiated this study because existing studies did not investigate intratracheal inoculation in (older) male ferrets. Shi *et al*. did perform an intratracheal infection in female ferrets and found viral RNA in the nasal turbinates and trachea, but not in the lung ^30^. In our study, adult male ferrets displayed higher viral titers after i.t. infection compared to young ferrets, indicating that age increases susceptibility to LRT SARS-CoV-2 infection in the ferret model. We postulated that the reduced viral replication in the LRT of especially young ferrets might be due to differential expression of the SARS-CoV-2 receptors ACE2 and TMPRSS2 ^1, 38^. However, this does not seem to be the case as ACE2 and TMPRSS2 expression was similar between nasal turbinate and lung tissue and young ferrets actually expressed more ACE2 and TMRPSS2 than adult ferrets. Combined with the observation that we did detect viral infection of the LRT by RT-qPCR for genomic RNA and subgenomic mRNA, it seems that factors other than receptor expression prevent the successful replication of SARS-CoV-2 in the LRT.

Notwithstanding the lack of clinical symptoms, we and others ^30, 32^ did find pathological abnormalities in infected ferrets. While SARS-CoV-2 replicated less efficiently in young i.t. inoculated animals and almost no adaptive immune response was induced, they did display some of the most affected pathology. It is however important to mention that all young ferrets euthanized on 21 dpi tested positive for antibodies against ferret corona and NL63 prior to SARS-CoV-2 infection. We cannot exclude that this might have had an effect and it has been postulated that existing cross-reactive immune responses can worsen COVID-19 disease outcome ^50^. Ryan *et al.* also found that pathology was still increasing between 14 and 21 dpi, although they did not report the immune status of their ferrets for other corona viruses. It is tempting to speculate that the increasing pathology post SARS-CoV-2 infection partly resembles aspects of long-COVID that have been described in convalescent patients ^22, 51–53^. These patients suffer from sequala up to months after initial infection with SARS-CoV-2. Although the symptoms are diverse, dyspnea is a relatively common occurrence. In theory, the BALT hyperplasia that we observed in ferrets and the resulting constriction of the bronchi(oli) could induce dyspnea, although there is no evidence yet that this is the cause of dyspnea in patients suffering from long-COVID.

After SARS-CoV-2 infection we detected antibodies against SARS-CoV-2 S-protein and RBD. Antibody titers were highest in i.n. infected ferrets, low in adult i.t. infected animals and absent in most young i.t. infected animals. This suggest that sufficient viral replication is required for seroconversion, as ferrets with lower viral loads also displayed lower humoral responses. This was especially clear in the i.t. young group, where the only ferret that developed humoral and cellular immunity also possessed the highest titre of replicative competent virus. Cellular responses were also strongest in i.n. inoculated ferrets. Similar to cellular responses of convalescent patients ^43, 54^, most responses in ferrets were aimed against S, M and N proteins. In both ferrets and humans ^43, 54^, almost no response against the E protein was observed. Cellular responses against multiple accessory proteins of SARS-CoV-2 were limited in the blood, but lung-derived lymphocytes did respond against several of these peptide pools. Likely, the relative abundance of SARS-CoV-2 specific T cells is higher in the lung, thereby increasing the sensitivity of the ELISpot assay. As cross-reactive cellular responses for SARS-CoV-2 have been described in humans ^40–43^, we also measured responses in the blood against peptide pools of S-proteins of other corona viruses. However, responses were low and interpretation of the results will require a larger group size and more sensitive assays.

Due to the global circumstances at the time of this study, there are several caveats present in the experimental set-up. The availability of (male) ferrets was limited and hence we could not investigate all groups at every timepoint. In addition, several ferrets were previously exposed to Aleutian disease and coronaviruses other than SARS-CoV-2 (details in Supplemental data). Although we did not find evidence that this infection history influenced our results, we cannot fully exclude that possibility. Finally, due to the limited availability of animals it will be difficult to model SARS-CoV-2 infections in older ferrets. However, as we did not find strong differences between young and adult ferrets upon intranasal inoculation, the use of young ferrets should suffice for future experiments.

As has been discussed here, ferrets are readily infected with SARS-CoV-2 but do not present clinical symptoms. Infected ferrets might thus represent the asymptomatic COVID-19 that manifests in a significant part of the population ^55^. In addition, the derailed immune responses in the respiratory tract in ferrets might model the long-term respiratory effects observed in long-COVID patients, although more in-depth research is required to verify this. Lastly, with the recently developed reagents for humoral and cellular immunology in the ferret model ^56–60^, vaccine-induced immune responses can be quantified and their effect on viral replication and pathology can be measured.

This matured ferret model can help with improving our understanding of SARS-CoV-2, thereby driving the development of new therapies and vaccines.

## Materials & Methods

### Ethical statement

All animal experiments were conducted in line with EU legislation. The experiment was approved by the local Authority for Animal Welfare of the Antonie van Leeuwenhoek terrain (Bilthoven, The Netherlands) under permit number AVD3260020184765 of the Dutch Central Committee for Animal experiments. Animals received food and water ad libitum and were inspected daily. If animals reached the pre-defined end points, they would be euthanized by cardiac bleeding under anesthesia with ketamine (5 mg/kg; Alfasan) and medetomidine (0.1 mg/kg; Orion Pharma). Endpoints were scored based on clinical parameters for activity (0 = active; 1 = active when stimulated; 2 = inactive and 3 = lethargic) and impaired breathing (0 = normal; 1 = fast breathing; 2 = heavy/stomach breathing). Animals were euthanized when they reached score 3 on activity level (lethargic) or when the combined score of activity and impaired breathing reached 4.

### Cell and virus culture

VERO E6 cells were cultured in DMEM (Gibco; Thermo Fisher Scientific) supplemented with 1x penicillin-streptomycin-glutamine (Gibco) and 10% fetal bovine serum (FBS; HyClone, GE Healthcare). SARS-CoV-2 virus was originally isolated from a Dutch patient (hCoV-19/Netherlands/ZuidHolland_10004/2020) and grown on VERO E6 cells in infection medium consisting of DMEM, 1x penicillin-streptomycin-glutamine and either 0% or 2% FBS. When >90% cytopathic effect (CPE) was observed, the suspension was spun down for 10 minutes at 4000x g to pellet cell debris. The remaining suspension was aliquoted and stored at −80°C. Sequencing of virus stock used for infection, revealed that no mutations had occurred compared to the primary isolate (GISAID accession ID: EPI_ISL_454753) and that the furin cleavage site was maintained. Wild-type mumps virus (MuVi/Utrecht.NLD/40.10; genotype G) was grown on Vero cells in DMEM (Gibco) with 2% FBS. Upon >90% CPE, the supernatant of the infected Vero cells was centrifuged at 500x g, filtered (5um pore size) and aliquots were stored at −80°C.

### Animal handling

Young (9-10 months) and adult (36-48 months) outbred male ferrets (Schimmel b.v., The Netherlands) arrived at the animal research facility (ARC, Bilthoven, The Netherlands) at least one week before commencement of the study. From arrival till the day of infection, animals were housed in open cages per group. Animals with confirmed exposure to Aleutian disease or NL63 were housed in a separate chamber. Infections were carried out in BSL-3 classified isolators where the animals remained till the end of the experiment. Placement of temperature transponders and infections were carried out under anesthesia with ketamine (5 mg/kg) and medetomidine (0.1 mg/kg). Buprenodale (0.2ml; AST Farma) was administered after transponder placement as a post-operative analgesic. Anesthesia was antagonized with atipamezole (0.25 mg/kg; Orion Pharma), which was delayed by 30 minutes in case of infections to prevent secretion of the inoculum by sneezing or coughing. Weight determinations and swabbing occurred under anesthesia with ketamine alone and did not require an antagonist.

### Study design

The study was split into three experiments (A, B and C) that were infected separately with SARS-CoV-2. Animals in the A, B and C experiments were respectively dissected 5, 14 and 21 days after infection. Each experiment consisted of 4-5 groups with 3 animals each. Groups were defined by their age (young vs adult) and infection route (intra nasal vs intra tracheal). Due to a shortage of adult animals, experiment B did not contain a group with i.n. infected adult animals. An additional mock-infected group to control for immunological assays and pathology was added to experiments B (young animals) and C (adult animals).

On day 0, ferrets were infected i.t. or i.n. with 10^7^ TCID_50_ SARS-CoV-2 diluted in PBS. Mock-infected ferrets received PBS i.n. alone. Virus was administered in 0.1ml for i.n. and 3ml for i.t administration. Prior to infection and on every-other day starting from day 3, bodyweight was measured and nasal, throat and rectal swabs were collected. At the end of the experiment, animals were euthanized by heart puncture under anesthesia with ketamine and medetomidine. Prior to heart puncture, bodyweight was measured and swabs were taken.

Importantly, this study had several complicating factors. At the supplier, males were housed in groups of 3 animals. Due to strict hierarchy present in male ferret groups, we were unable to randomly allocate individual animals to treatment groups. Instead, whole cages (with 3 animals) were randomly allocated to a certain treatment. Before arrival at the animal facility, ferret sera were screened by the European Veterinary Laboratory (EVL, The Netherlands) for prior infections with Aleutian disease, human corona NL63, canine distemper virus (CDV), feline corona virus (FCoV) and canine enteric corona virus (CCoV). FCoV and CCoV are representative for systemic and enteric ferret corona viruses respectively. Almost all animals displayed prior exposure against CDV and multiple animals tested positive for antibodies against NL63, FCoV and CCoV before start of the experiment (Supplementary table 1).

### Sample collection

During the experiment, blood was collected from the vena cava and stored in either sodium-heparin coated VACUETTE tubes (Greiner) for cellular assays or in CAT serum separator clot activator VACUETTE tubes (Greiner) to isolate serum. Blood was kept at RT and analyzed the same day. At the end of the experiment, blood was collected by heart puncture after which groups B and C received a lung perfusion to remove the majority of lymphocytes in the circulation that might affect cellular assays. The lower 4cm of the trachea was stored in formalin to study pathology while the middle 1.5cm was used to determine viral load. Lungs were weighed and the left cranial lobe was inflated with formalin and stored in 10% buffered formalin for histopathological analysis. To assess viral RNA and TCID_50_-titers, 0.5cm slices of the right cranial, middle and caudal lobe were collected in Lysing Matrix A tubes (MP Biomedicals, Germany). Samples in Lysing Matrix A tubes were stored at −80°C until analysis. The rest of the lungs were used for immunological analysis and were kept cold (4°C) o/n until processing the next day.

Of the intestine, 1.5cm parts of the ileum and upper colon were collected in Lysing Matrix A tubes and formalin to assess viral RNA and pathology respectively. Next, the cranium was bisected with an oscillating blade moving from the caudal to the rostral position to prevent contamination of brain tissue by any particles from URT. The nasal turbinates on the left of the septum were used for pathology while the right side was used for virology and immunology. Finally, the olfactory bulb (OB) and sections of the cerebrum and cerebellum were collected for virology and pathology. Samples for pathology, virology and immunology were stored respectively in formalin, Lysing Matrix A tubes and RPMI medium supplemented with 1x penicillin-streptomycin-glutamine.

Nasal and throat swabs were collected in 2ml transport buffer consisting of 15% sucrose (Merck), 2.5μg/ml Amphotericin B, 100 U/ml penicillin, 100μg/ml streptomycin and 250μg/ml gentamicin (all from Sigma). Rectal swabs were stored in 1ml S.T.A.R. buffer (Roche). After collection, all swabs were vortexed, aliquoted under BSL-3 conditions and stored at −80 °C until further analysis. Of each sample, 200μl was directly added to MagNA Pure External Lysis Buffer (Hoffmann-La Roche, Basel, Switzerland), vortexed and stored at −20 °C for RT-qPCR. Samples stored in Matrix A tubes were thawed and 750μl of DMEM infection medium (DMEM containing 2% FBS and 1x penicillin-streptomycin-glutamine) was added. Tissues were then dissociated in a FastPrep-24™ by shaking twice for 1 minute after which the samples were spun down for 5 minutes at 4000x g. Of the supernatant, 200μl was used for RT-qPCR analysis as detailed above and 250μl was used for TCID_50_-determination.

The dissection of SARS-CoV-2 infected animals occurred under BSL-3 conditions and all materials from swabs, samples in Lysing Matrix A tubes and nasal turbinates were handled under BSL-3 conditions. Blood was handled under BSL-2 conditions as blood was shown to be PCR-negative for SARS-Cov-2. Spleen and lung dissected 14 and 21 dpi were processed under BSL-2+ conditions as lung tissue did not contain infectious virus 5 d.p.i. as shown by TCID50 analysis.

### Temperature logging

In experiments B and C, animals received temperature probes (Star Oddi, Iceland) two weeks before the infection. These probes recorded temperature every 30 minutes from 7 days before infection till the end of the experiment. Fever was calculated as deviation from baseline (ΔT), where the baseline refers to the mean temperature over 5 days prior to SARS-CoV-2 infection.

### Pathology

Pathology scoring was performed as described before [16, 28]. After fixation, the lung lobes were embedded in paraffin and sliced into 5μm thick sections. Slides were stained with haematoxylin and eosin and microscopically examined at 50x or 100x magnification. For each tissue, at least 20 microscopic fields were scored. Pathological scoring distinguished between the categories ‘epithelial damage’ and ‘inflammation’. Damage related parameters included hypertrophy, hyperplasia, flattened or pseudo squamous epithelia, necrosis and denudation of bronchi(oli) epithelium, hyperemia of septa and alveolar emphysema and haemorrhages. Inflammation related parameters included (peri)bronchi(oli)tis, interstitial infiltrate, alveolitis and (peri)vasculitis characterized by polymorphonuclear (PMN) cells, macrophages and lymphocytic infiltrate. Pathological findings were scored on a scale of 0 (no aberrations) to 5 (severe damage) and were summarized in two ‘end scores’ for the categories ‘epithelial damage’ and ‘inflammation’. Microscopic slides were randomized and scored blindly by an experienced pathologist.

### Lymphocyte isolation

Blood collected in sodium-heparin tubes was diluted 1:1 with PBS (Gibco) and layered on top of a 1:1 mixture of Lymphoprep (1.077 g/ml, Stemcell) and Lymphoyte-M (1.0875 g/ml, Cedarlane). The gradient was spun at 800x g for 30 minutes and the interface containing PBMCs was collected. The cells were subsequently washed twice with washing medium (RPMI1640 + 1% FBS) and spun down at 500x g for 10 minutes (first wash) or 5 minutes (second wash). After the final washing step, cells were resuspended in stimulation medium (RPMI1640 + 10% FBS + 1x penicillin-streptomycin-glutamine).

On days 14 and 21 after infection, the lungs of euthanized animals were perfused with saline to remove circulating lymphocytes from the lungs as described before ^56^. Lymphocytes from the lung were isolated using enzymatic digestion. Lungs were first processed into small dices of approximately 5mm^3^ using scissors. The diced tissue was then digested for 60 minutes at 37°C in 12ml of a pre-heated suspension of collagenase I (2.4mg/ml, Merck) and DNase I (1mg/ml, Novus Biologicals) in RPMI1640 while rotating. Tissue was further homogenized by gently pressing the tissue over a sieve using the plunger of a 10ml syringe. The resulting suspension was diluted with EDTA-supplemented washing medium (RPMI1640 + 1% FBS + 2mM EDTA (Invitrogen)) and filtered over a 70μm cell strainer. This suspension was layered on top of 15ml Lympholyte-M in a 50ml tube. Density centrifugation and washing steps were performed as described above, with the exception that washing medium was supplemented with EDTA to prevent agglutination of cells

### SARS-CoV-2 peptide pools

PepMix™ peptide pools for T cell stimulation assays were obtained from JPT Peptide Technologies GmbH. Each pool contained 15 amino acids long peptides with an overlap of 11 amino acids spanning an entire protein of SARS-CoV-2. Due to the length of the spike protein, the spike PepMix™ was distributed over two separate vials containing peptides 1-158 and 159-315.

### Elispot

Pre-coated Ferret IFNγ-ELISpot (ALP) plates (Mabtech) were used according to the manufacturers protocol. Per well, 250K PBMCs or 31.25K lung lymphocytes were stimulated with live virus (MOI 1) or SARS-CoV-2 peptide pools (1ug/ml) in Elispot plates for 20 hours at 37°C. Plates were then washed and developed according to the manufacturers protocol, with the modification that incubation with the first antibody occurred o/n at 4°C instead of 2 hours at RT. After the final washing step, plates were left to dry for >2 days. Plates were then packaged under BSL-3 conditions and heated to 65°C for 3 hours to inactivate any remaining SARS-CoV-2 particles. Plates were analyzed on the ImmunoSpot^®^ S6 CORE (CTL, Cleveland, OH). Spot counts were corrected for background signals by subtracting the number of spots in the medium condition from all other conditions. Data were visualized on a log-scale, so the minimum spot count was set to ‘1’ for visualization purposes.

### Trucount

Of each animal, 50μl of heparin-blood was used for trucount analysis with the non-centrifugation PerFix-NC kit (Beckman Coulter). Cells were first stained extracellular with α-CD4-APC (02, Sino Biological), α-CD8a-eFluor450 (OKT8, eBioscience), and α-CD14-PE (Tük4; Thermo Fisher) for 30 minutes at RT. Cells were then fixated with 5μl Fixative Reagent for 15 minutes after which 300μl Permeabilizing reagent was added. The subsequent intracellular staining consisted of α-CD3e-FITC (CD3-12, Biorad) and α-CD79a-APC/eFluor780 (eBioscience). After 30 minutes incubation at RT, 3ml of Final reagent was added to each sample. To decrease measurement time, samples were spun down for 5 minutes at 500x g and 2.8ml suspension was removed. The pellet was resuspended in the remaining volume and 50μl of Precision Count beads (Biolegend) was added to each sample to calculate the absolute number of cells. Samples were measured on a FACSymphony A3 (BD) and analysed using FlowJo™ Software V10.6.2 (BD). An example of the gating strategy is present in Supplemental figure 1.

### Virus titer analysis

Virus stocks were titrated in 8-plo on VERO E6 cells in 96-wells plates. Samples were titrated in DMEM medium containing 2% FBS and 1x penicillin-streptomycin-glutamine. After 6 days, CPE was scored and TCID_50_ values were calculated using the Reed & Muench method. Nose and throat swabs were similarly tested, but in 6-plo with 8 dilutions. For swabs, 2.5μg/ml Amphotericin B and 250μg/ml gentamicin was added to the titration medium.

### RT-qPCR

Lysis buffer was spiked with equine arteritis virus (EAV) as an internal RT-qPCR control and stored at −20°C until sample material was added. Total nucleic acid was extracted from samples with the MagNA Pure 96 system (Hoffmann-La Roche) using the MagNA Pure 96 DNA and Viral NA Small Volume Kit and eluted in a volume of 50μl Roche Tris-HCl elution buffer. A 20μl Real-time Reverse-Transcription PCR (RT-qPCR) reaction contained 5μl of sample nucleic acid, 7μl of 4x Taqman Fast Virus Master Mix (Thermo Fisher), 5μl of DNAse/RNAse free water and 3μl of primers and probe mix (sequences shown in Table 1). The in-house SARS-CoV-2 detection assay was performed using E-gene primers and probe specific for SARS-related betacoronaviruses as described by Corman et al. ^61^ and the equine arteritis virus (EAV) internal control primers and probe as described by Scheltinga et al. ^62^. This E-gene RT-qPCR detects genomic and subgenomic SARS-CoV-2 RNA molecules. The in-house subgenomic mRNA E-gene assay was performed using the E-gene reverse primer and probe and the forward primer as described by Zhang et al. ^63^. All tests were performed on a Light Cycler 480 I (LC480 I, Roche) according to the cycling protocol detailed in table 2. Cycle threshold (Ct) values were recorded.

**Table 1.**
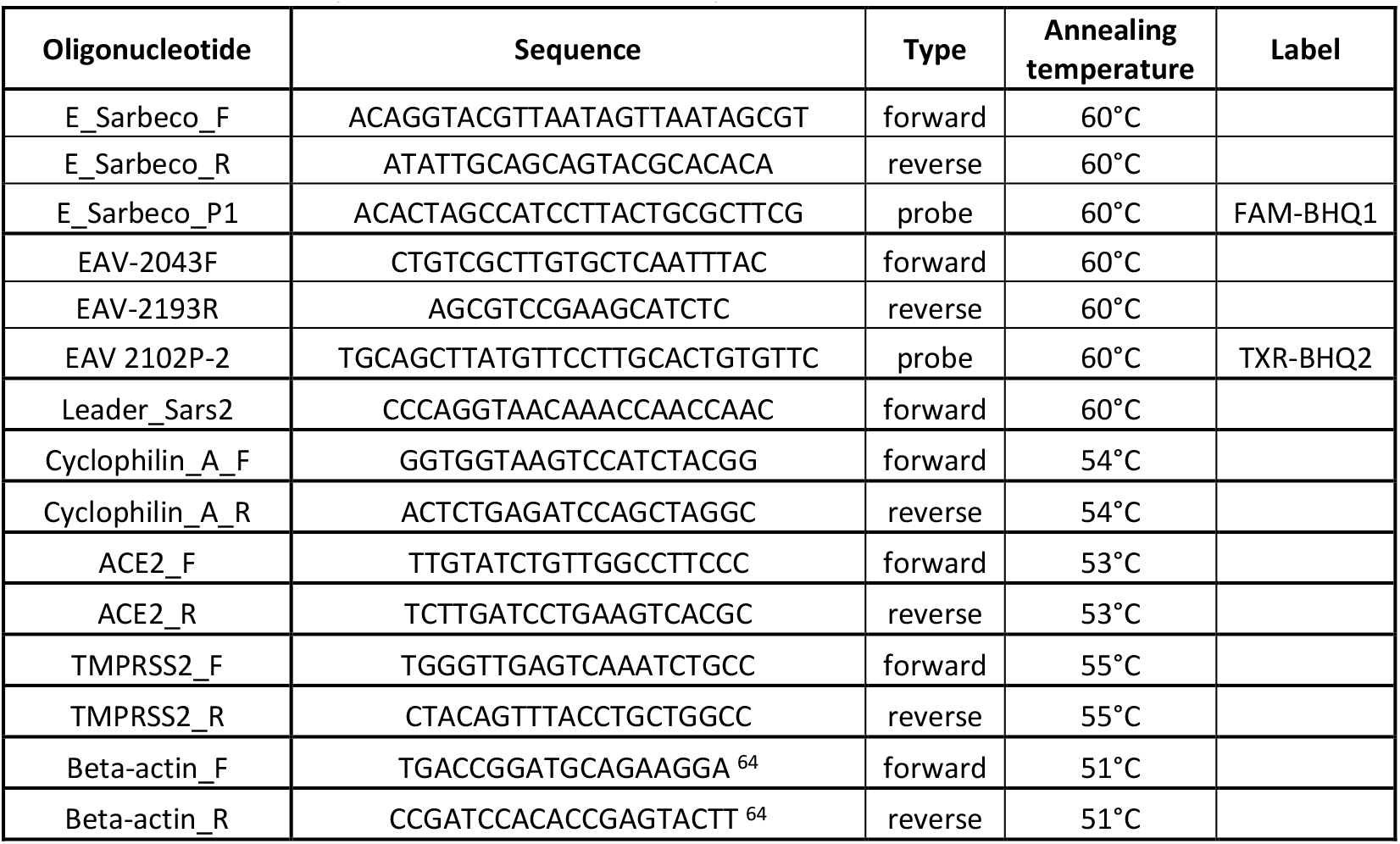
Primers and probes used in this study

**Table 2.**
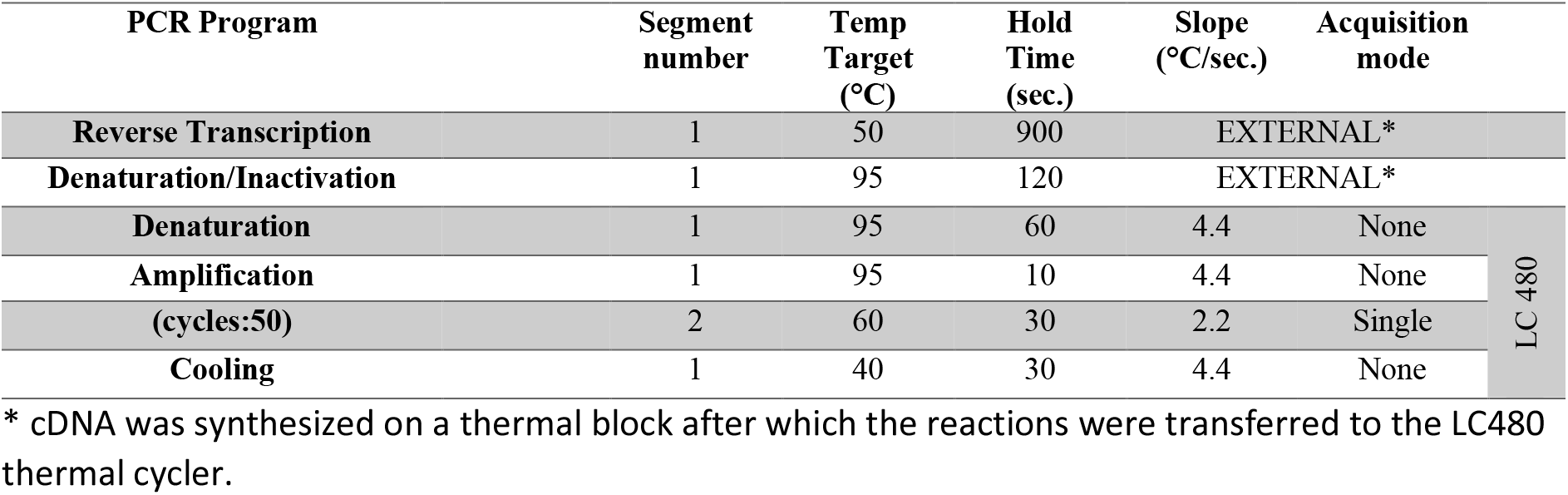
Cycling and temperature protocol for SARS-CoV-2 E-gene RT-qPCR

Total nucleic acid extracted with the MagNA Pure 96 system was used to determine ACE2 and TMPRSS2 expression in a separate reaction. First, cDNA was synthesized from the isolated RNA with the iScript™ cDNA synthesis kit (Bio-Rad) according to the manufacturers protocol using a StepOnePlus RT-PCR system (Thermo Fisher). RT-qPCR was performed using 1x Maxima SYBR Green/ROX qPCR Master Mix (Thermo Fisher) on the StepOnePlus with primers for Cyclophilin A, ACE2, TMPRSS2 and Beta-actin (Isogen Life Science, Netherlands; Table 1). RT-qPCR was performed for 10 minutes at 95°C, followed by 40 cycles of 95°C for 30 seconds and primer-specific annealing temperatures (see above) for 45 seconds. This was followed by 95°C for 15 seconds, 53°C for 60 seconds and 95°C for 15 seconds. A pooled reference sample consisting of 5-fold dilutions of cDNA of the nasal turbinates of 6 animals was taken along in duplicate with each RT-qPCR to create a standard curve. RNA concentrations (in arbitrary units) of test samples were interpolated from the standard curve. The relative mRNA expression of target genes ACE2 and TMPRSS2 was calculated by dividing the interpolated arbitrary RNA unit of the target genes by the geometric mean of the endogenous control genes (Cyclophilin A and Beta-actin).

### ELISA

Immulon 2 HB 96-well plates (Thermo Fisher) were coated overnight at RT with 100μl/well 0.25μg/ ml recombinant SARS-CoV-2 Spike or Spike receptor binding domain protein (Sino biological, China) and washed thrice with PBS + 0.1% Tween-80 before use. Sera were first diluted 1:100 in PBS + 0.1% Tween-80 and then 2-fold serially diluted. Per well, 100μl of diluted sera was added and plates were incubated for 60 minutes at 37°C. After washing thrice with 0.1% Tween-80, plates were incubated for 60 minutes at 37°C with HRP-conjugated goat anti-ferret IgG (Alpha Diagnostic), diluted 1:5000 in PBS containing 0.1% Tween-80 and 0.5% Protivar (Nutricia). Plates were then washed trice with PBS + 0.1% Tween-80 and once with PBS, followed by development with 100μl SureBlue™ TMB (KPL) substrate. Development was stopped after 10 minutes by addition of 100μl 2M H_2_SO_4_ and OD_450_-values were determined on the EL808 absorbance reader (Bio-Tek Instruments). Individual curves were visualized using local polynomial regression fitting with R software ^65^. Antibody titers were determined as the dilution at which antibody responses dropped below background. This background was calculated as the ‘mean + 3 * standard deviation’ of the OD_450_ at a 400x serum-dilution of all animals tested before SARS-CoV-2 infection.

## Data analysis & Statistics

All raw data was analyzed with the software detailed above. These data were then exported to Excel and loaded into R software ^65^. Data analysis and visualization of data was carried out using the `R` packages `ggplot2` ^66^, `tidyverse` ^67^, `ggpubr` ^68^. Due to the exploratory goal and small group numbers of this study, statistical analysis was limited. Blood cell counts in young and adult animals was compared in `R` using the Wilcoxon signed-rank test and corrected for multiple testing by the Holm-Bonferroni method ^69^.

## Data availability

All raw data files are being stored in-house on backed-up servers and are available upon reasonable request to the corresponding author. The sequence of the SARS-CoV-2 isolate can be found on GISAID (accession ID: EPI_ISL_454753).

## Acknowledgements

We would like to thank Gabriel Goderski for isolating and growing the initial SARS-CoV-2 isolate and Jeroen Cremer for sequencing. We are also grateful to the biotechnicians from the animal facility for excellent care-taking of the animals and dr. Luytjes for critical reviewing of the manuscript.

## Author contributions

K.v.d.V., H.v.D., L.W., A.G., T.S., J.K. and S.L. performed experiments. K.v.d.V., P.R., P.K. and J.d.J. analyzed data. K.v.d.V. and J.d.J. designed the experiment and wrote the manuscript together with P.R., A.M. and P.K.

## Materials & Correspondence

Correspondence and requests for materials should be addressed to J.d.J.

---

**Supplemental Table 1.**
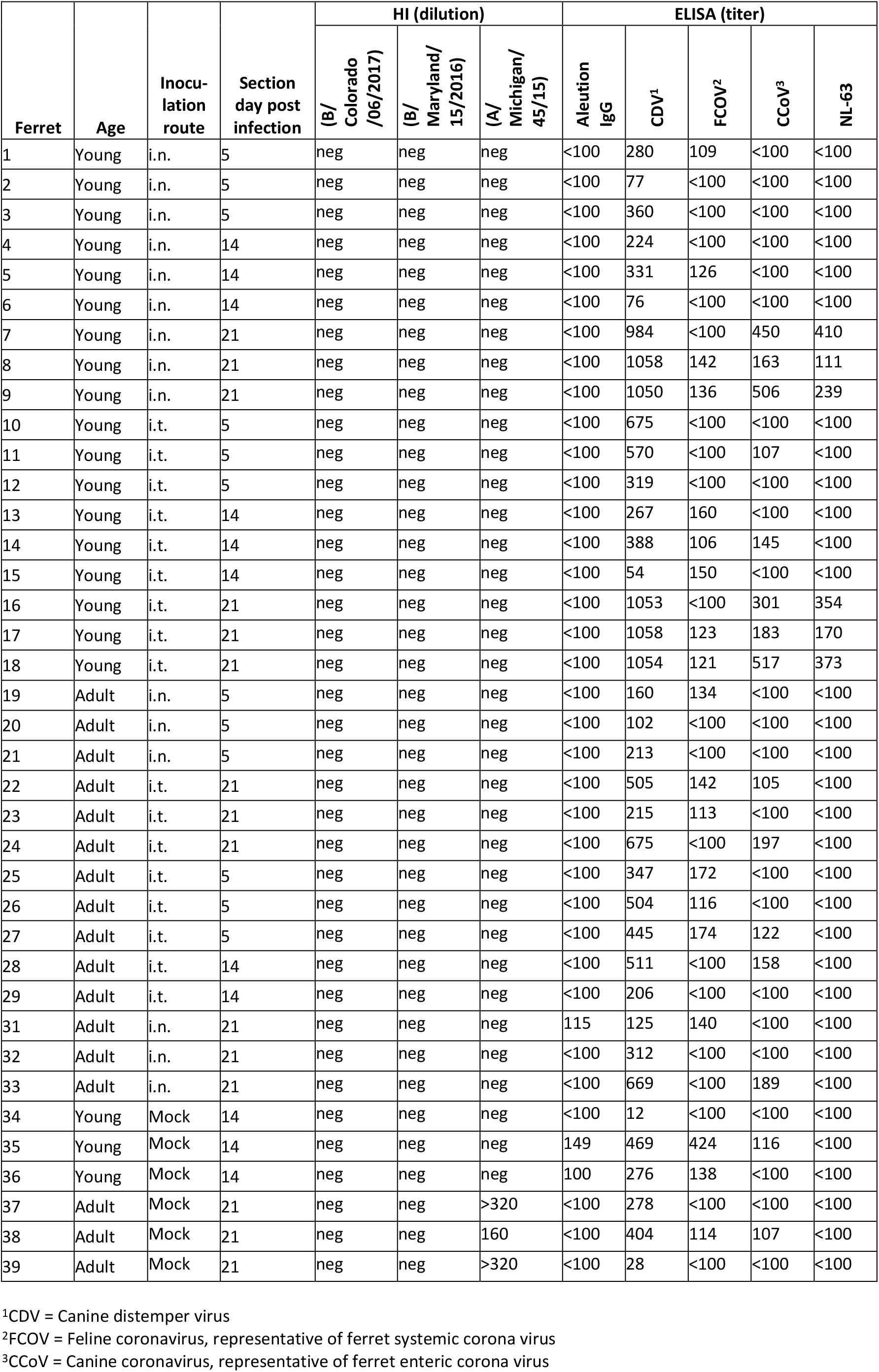
Immune status before SARS-CoV-2 infection.

**Supplemental figure 1:**
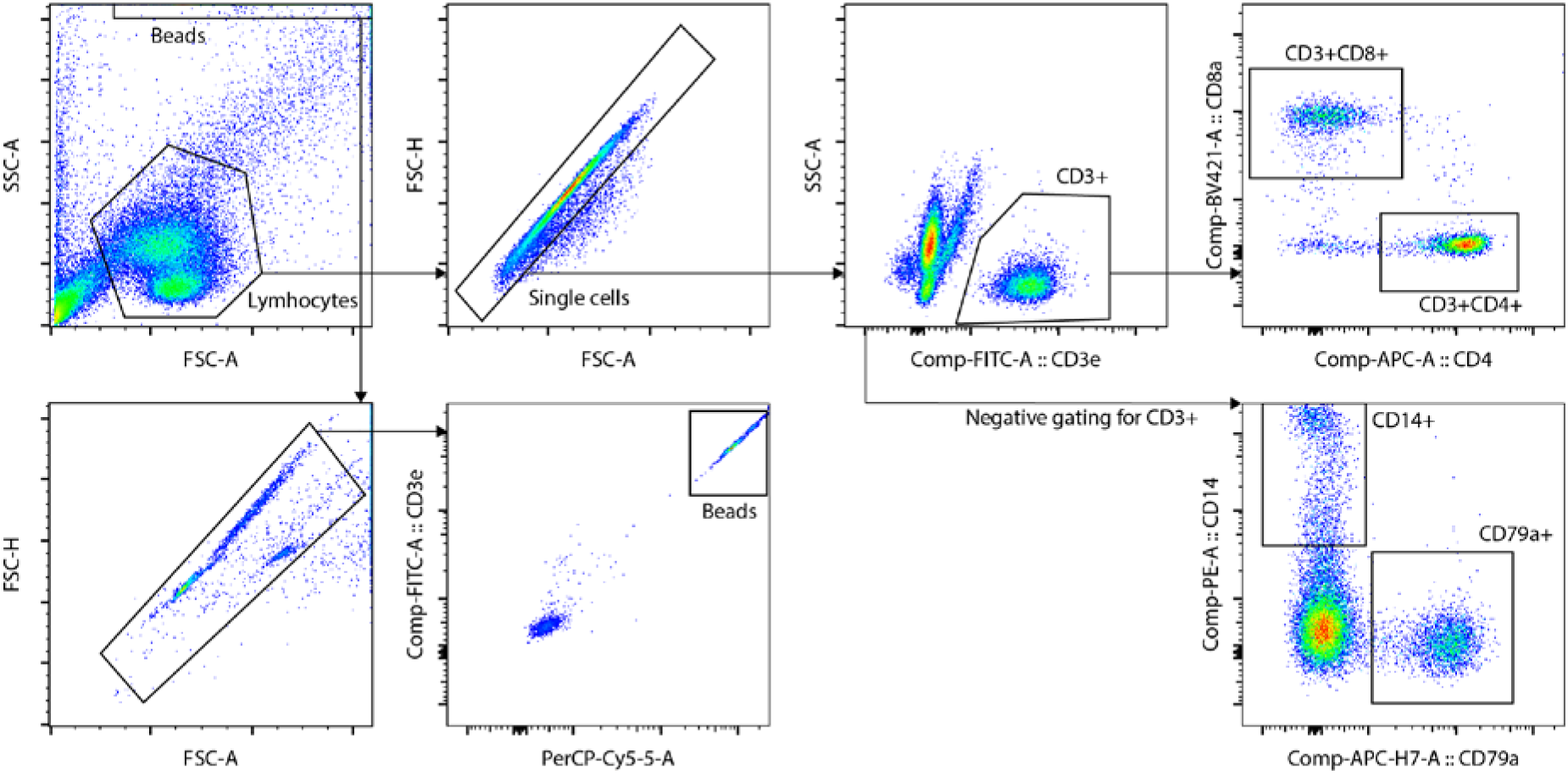
Gating strategy of whole blood trucount. Plots show the gating strategy for identification of cell subsets in whole blood of young and adult ferrets. CD14 and CD79a subsets were gated in the ‘Single cells’ population excluding all CD3+ cells. Beads were used to correct cell counts for the volume of measured blood.

**Supplemental figure 2:**
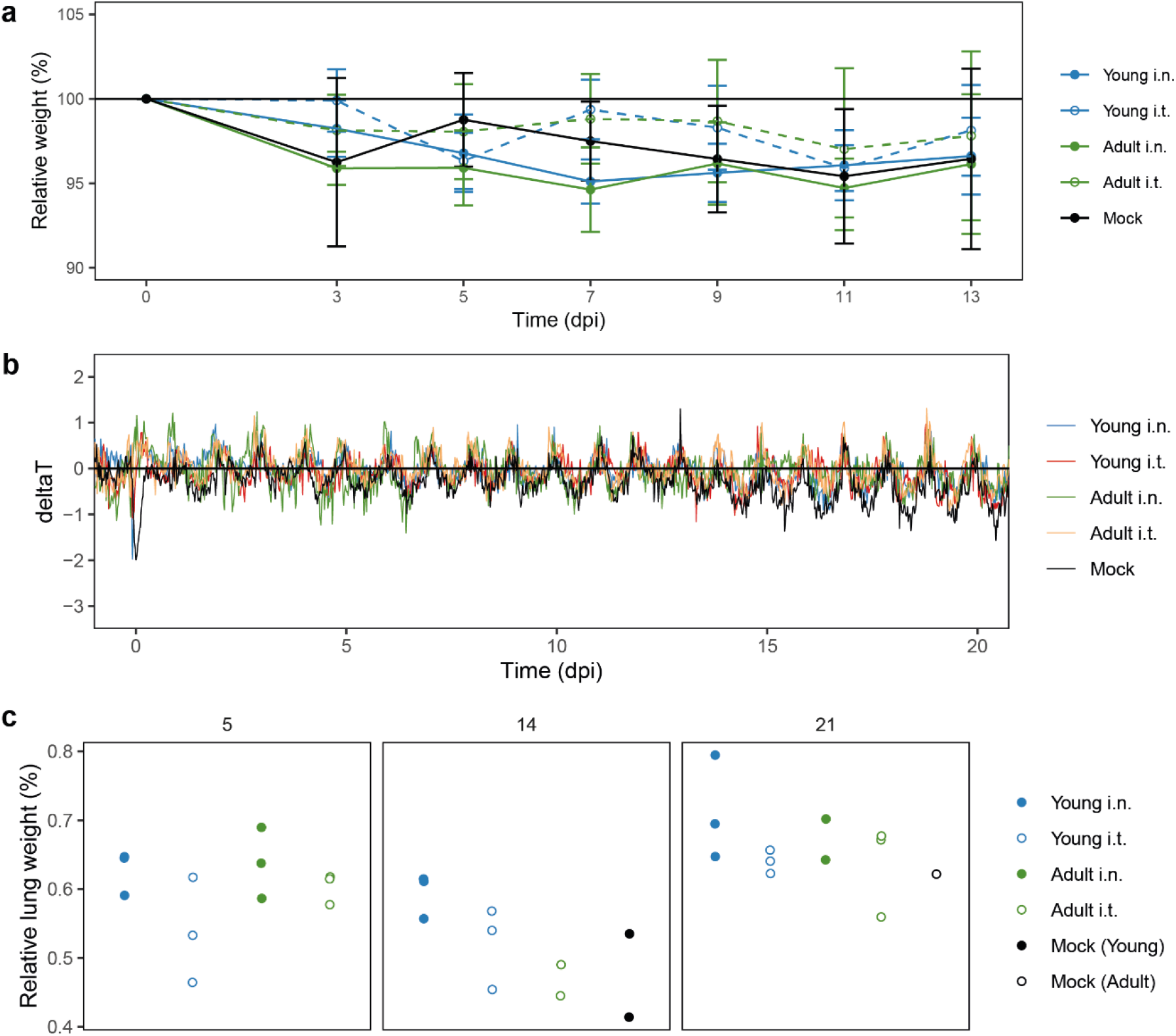
SARS-CoV-2 infection does not induce clinical disease in male ferrets. **A**) body weight was measured on various time points and depicted as % of original bodyweight on the day of infection. **B**) Body temperature was measured continuously in 30-minute intervals by implanted abdominal transponders. The ΔT was calculated by subtracting body temperature during baseline (1-6 days before infection) from the body temperature after SARS-CoV-2 infection. **C**) Relative lung weight depicted as a percentage of total body weight on the day of infection. The different panels depict the relative lung weight on 5, 14 and 21 days post infection (dpi). Lines (a,b) depict the group mean while the error bars depict standard error of the mean (a only). For a,b: n = 3-6; for c: with exception of ‘Adult i.t.’ on 14 dpi (n = 2), all groups are n = 3.

**Supplemental figure 3:**
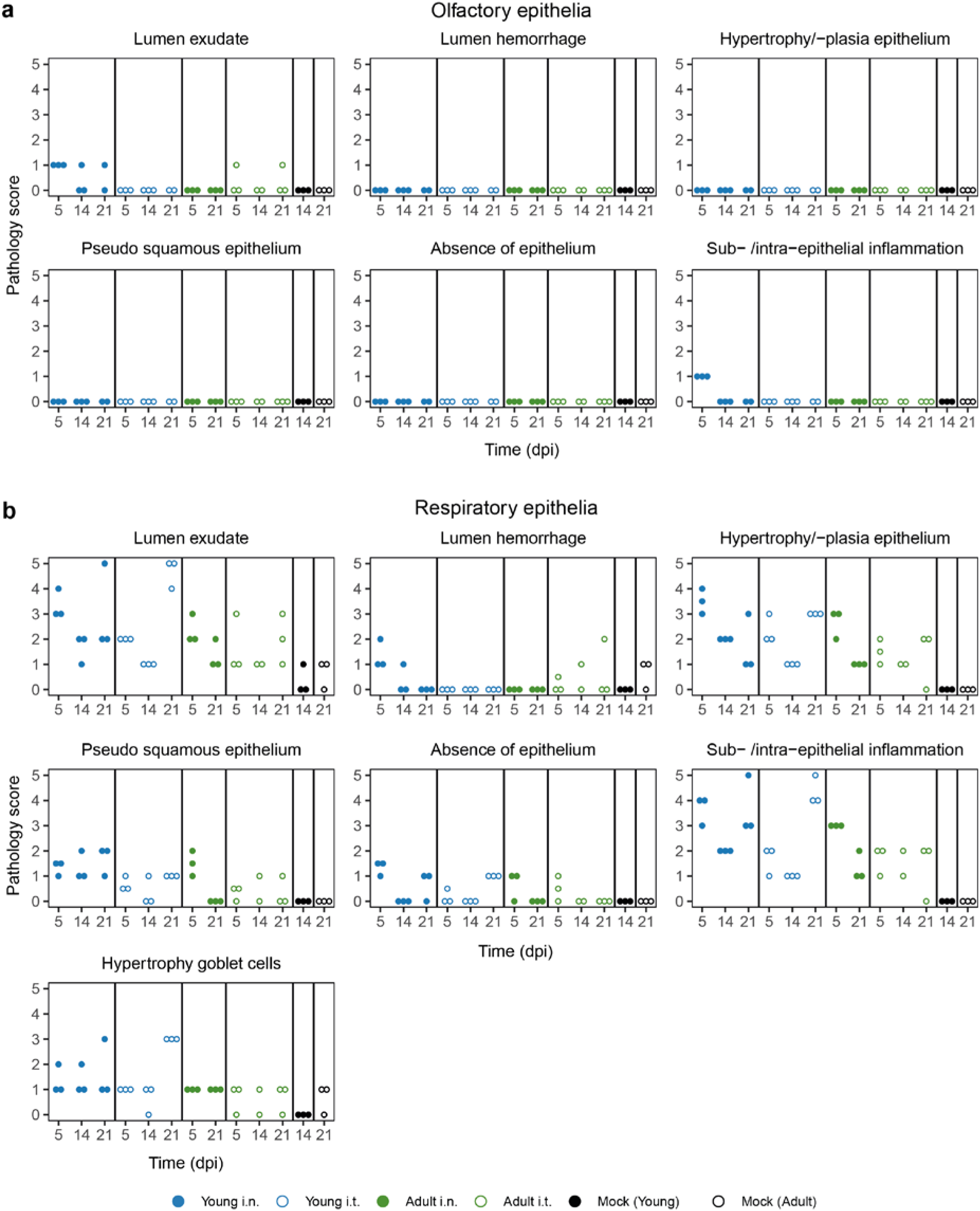
Extensive pathology scoring nasal turbinates. **A, B)**Scoring of (a) olfactory and (b) respiratory turbinates. Panels depict individual parameters related to epithelial damage and inflammation on 5, 14 and 21 days post infection (dpi). The infection-induced pathology was scored on a scale of 0–5 based on the parameters described in the materials and methods. With exception of ‘Adult i.t.’ on 14 dpi (n = 2), all groups are n = 3.

**Supplemental figure 4:**
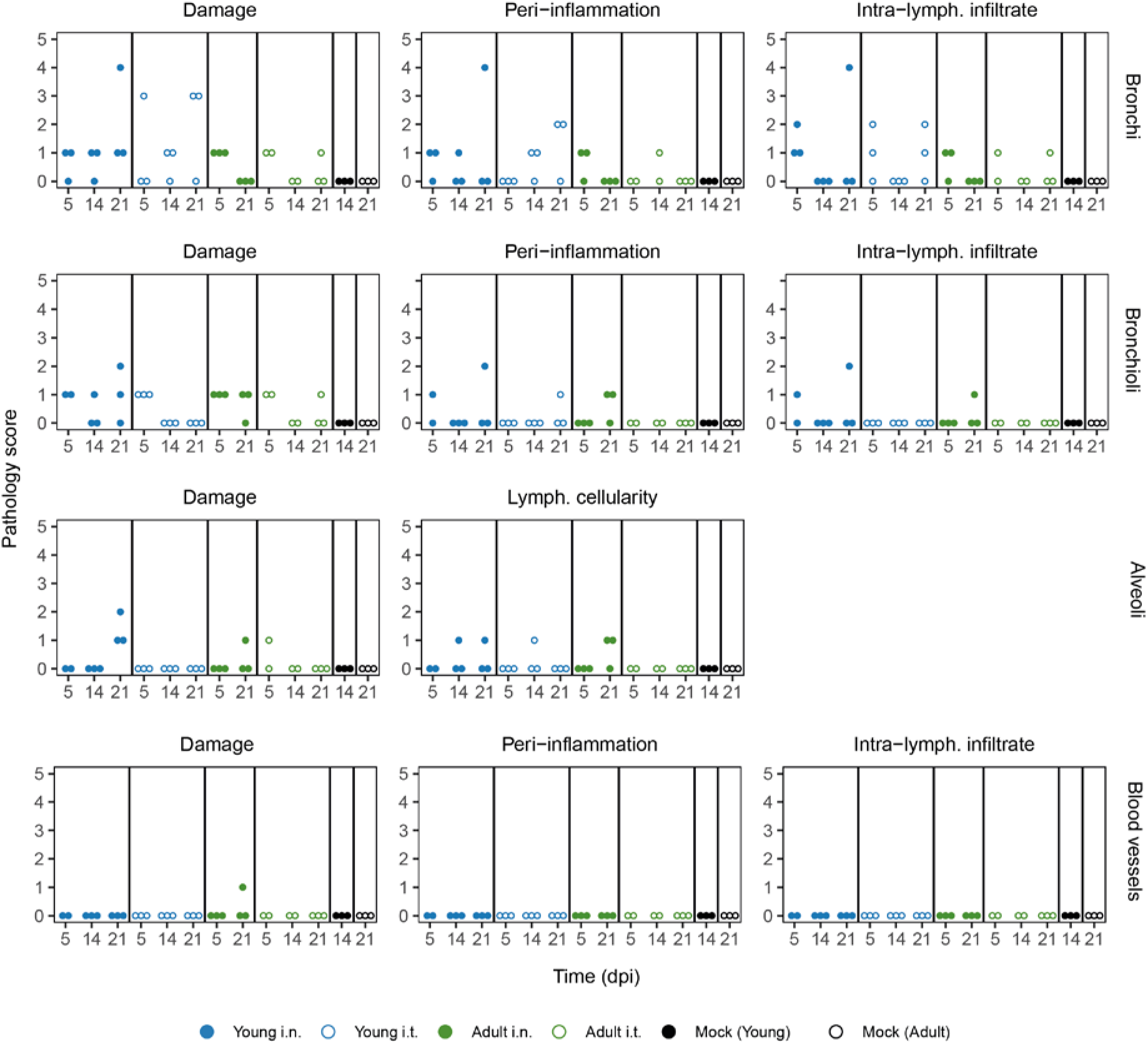
Extensive pathology scoring lungs. Panels show individual scoring by parameters related to epithelial damage and inflammation. The infection-induced pathology was scored on a scale of 0–5 based on the parameters described in the materials and methods on 5, 14 and 21 days post infection (dpi). With exception of ‘Adult i.t.’ on 14 dpi (n = 2), all groups are n = 3.

**Supplemental figure 5:**
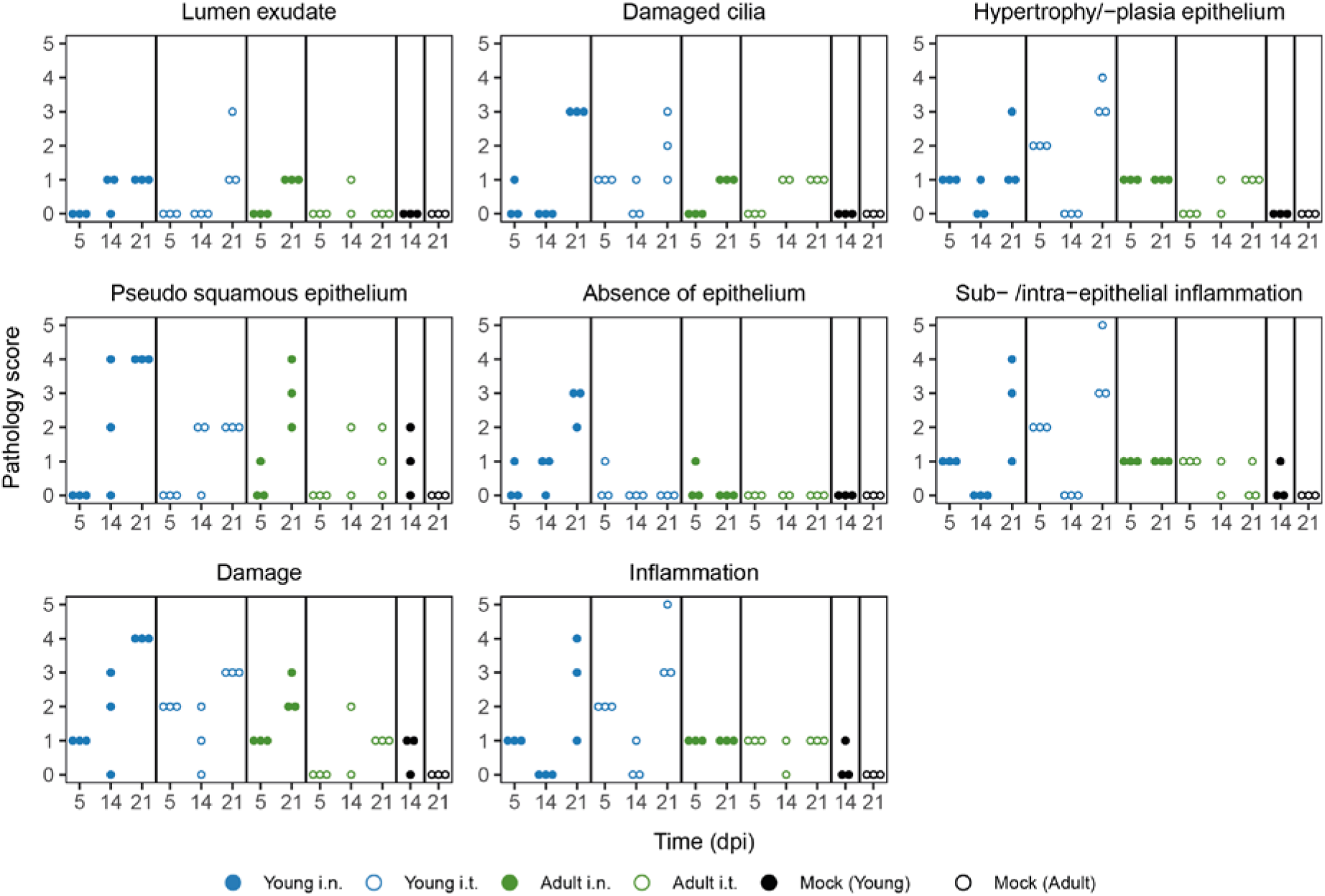
Extensive pathology scoring trachea. Panels show individual scoring by parameters related to epithelial damage and inflammation. The infection-induced pathology was scored on a scale of 0–5 based on the parameters described in the materials and methods on 5, 14 and 21 days post infection (dpi). With exception of ‘Adult i.t.’ on 14 dpi (n = 2), all groups are n = 3.

**Supplemental figure 6:**
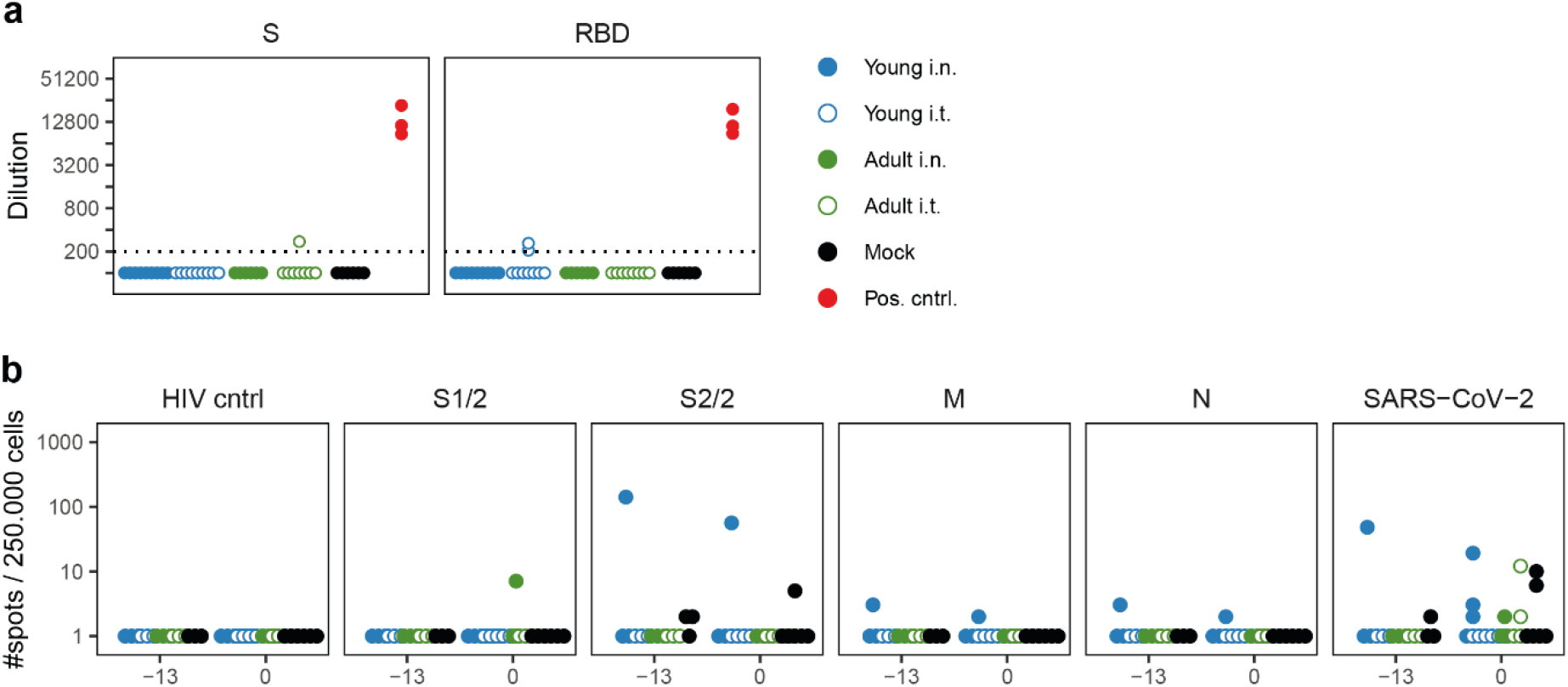
Pre-existing immune responses against SARS-CoV-2 in ferrets. **A**) Sera from three animals before SARS-CoV-2 infection contained small antibody responses against (the receptor binding domain [RBD] of) spike (S) in ELISA assays, but these were different animals from the one that responded in the ELISpot of panel B. Responses are depicted as the (modelled) dilution at which the ELISA curve drops below background (mean + 3x SD of SARS-CoV-2 naïve animals at 200x dilution). The dotted line indicates the lowest dilution tested and negative samples were set to half that dilution for visualization purposes. Positive control consists of sera of SARS-CoV-2 infected animals collected 21 days post infection (dpi). N= 3 for positive control sera and N = 6-9 for other groups **B**) IFNγ-ELISpots performed with PBMCs isolated 13 and 0 days before SARS-CoV-2 infection indicate that one animal already possessed (cross-reactive) T cell responses against overlapping peptide pools of the S-protein of SARS-CoV-2. Data shown were corrected for medium background and were set to a minimum of 1 spot for visualization on a log-scale. N = 3 at −13 dpi and n = 5-6 for 0 dpi.

**Supplemental figure 7:**
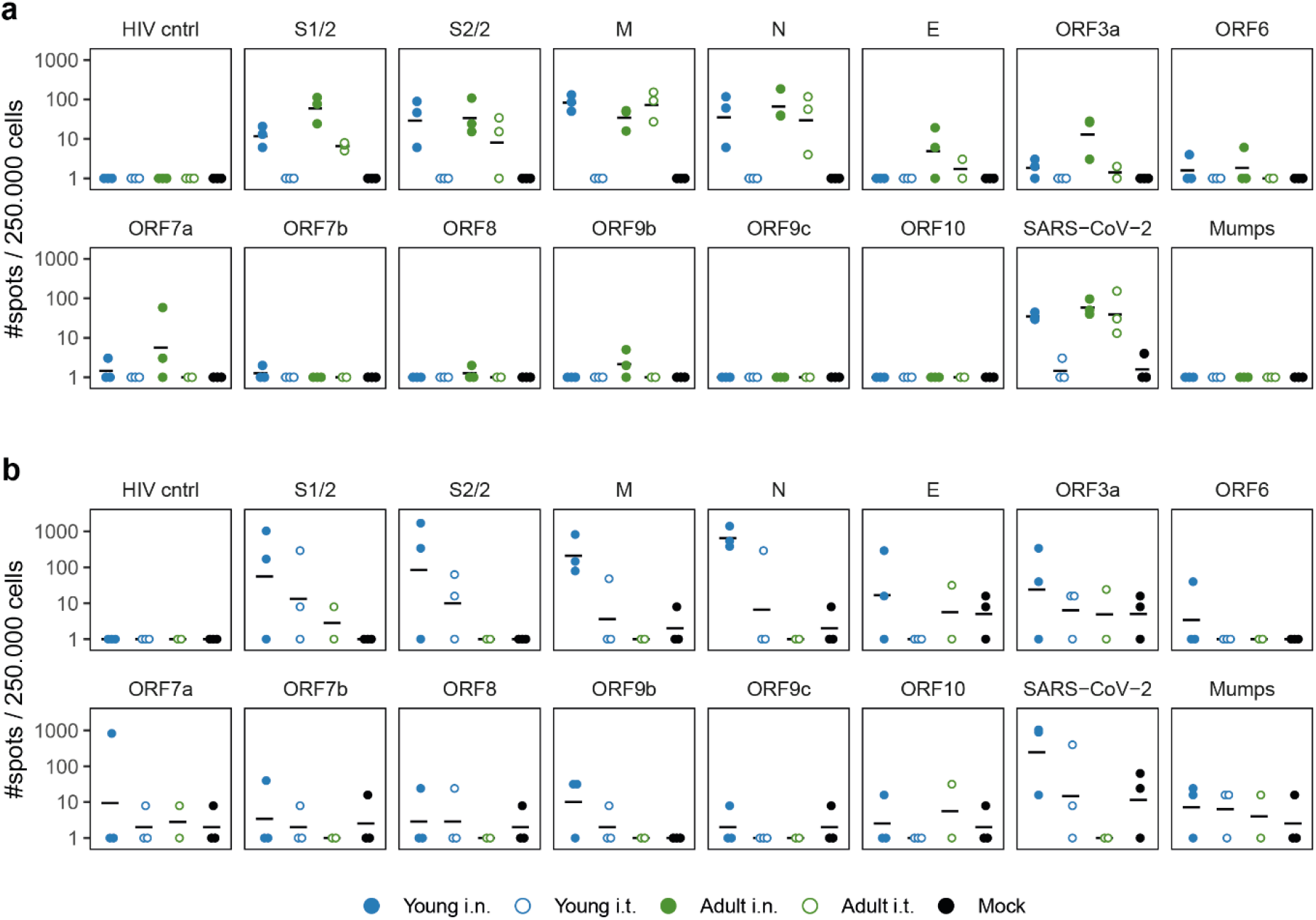
Cellular responses against SARS-CoV-2 in PBMC and lung. **A, B**) Cellular responses in PBMCs (a) and lung derived lymphocytes (b) as determined by IFNγ-ELISpot. Cells were stimulated with various SARS-CoV-2 peptide pools or live virus. Dots show individual ferrets while black lines indicate the group geometric mean. Data were corrected for medium background and were set to a minimum of 1 spot for visualization on a log-scale. **A**) Responses in PBMC isolated 21 days post infection (dpi). **B**) Responses of lung-derived lymphocytes 14 dpi. N = 3 for all panels, with exception of ‘Adult i.t.’ on 14 dpi (n = 2).

